# Distinct subpopulations of D1 medium spiny neurons exhibit unique transcriptional responsiveness to cocaine

**DOI:** 10.1101/2023.01.12.523845

**Authors:** Robert A. Phillips, Jennifer J. Tuscher, N. Dalton Fitzgerald, Ethan Wan, Morgan E. Zipperly, Corey G. Duke, Lara Ianov, Jeremy J. Day

**Author notes:** Correspondence to Jeremy Day* ( | day-lab.org | <u>@DayLabUAB</u>).

## Abstract

Drugs of abuse increase extracellular concentrations of dopamine in the nucleus accumbens (NAc), resulting in transcriptional alterations that drive long-lasting cellular and behavioral adaptations. While decades of research have focused on the transcriptional mechanisms by which drugs of abuse influence neuronal physiology and function, few studies have comprehensively defined NAc cell type heterogeneity in transcriptional responses to drugs of abuse. Here, we used single nucleus RNA-seq (snRNA-seq) to characterize the transcriptome of over 39,000 NAc cells from male and female adult Sprague-Dawley rats following acute or repeated cocaine experience. This dataset identified 16 transcriptionally distinct cell populations, including two populations of medium spiny neurons (MSNs) that express the Drd1 dopamine receptor (D1-MSNs). Critically, while both populations expressed classic marker genes of D1-MSNs, only one population exhibited a robust transcriptional response to cocaine. Validation of population-selective transcripts using RNA in situ hybridization revealed distinct spatial compartmentalization of these D1-MSN populations within the NAc. Finally, analysis of published NAc snRNA-seq datasets from non-human primates and humans demonstrated conservation of MSN subtypes across rat and higher order mammals, and further highlighted cell type-specific transcriptional differences across the NAc and broader striatum. These results highlight the utility in using snRNA-seq to characterize both cell type heterogeneity and cell type-specific responses to cocaine and provides a useful resource for cross-species comparisons of NAc cell composition.

## INTRODUCTION

Drug addiction is a chronic, complex neuropsychiatric disorder characterized by a loss of control of drug-taking behaviors. The current drug addiction and overdose epidemic in the United States has been worsened by the COVID-19 pandemic as people struggle with social isolation and economic distress^1^. Much of the current addiction epidemic and surge in overdose deaths has been attributed to the use of opioids and, in particular, synthetic opioids such as fentanyl. However, drug overdose deaths associated with psychostimulants rose 50% between 2019-2020^2^. Furthermore, drug overdose deaths associated with cocaine have increased 3-fold since 1999^2,3^. While pharmacological treatments exist for opioid use disorder, no such treatments are available for stimulant use disorder. Thus, continued research on the molecular adaptations that occur following exposure to cocaine, and the cell types that may be susceptible to these adaptations, is important for development of future therapeutics.

The mesolimbic dopamine pathway is heavily implicated in molecular, cellular, and behavioral responses to drugs of abuse. This pathway consists of dopaminergic neurons in the ventral tegmental area (VTA) that project to the nucleus accumbens (NAc). Dopaminergic neurotransmission in the NAc is important for adaptive behavior, as dopamine (DA) release tracks reward-predictive cues as a function of learning^4^. Exposure to drugs of abuse, such as cocaine, increases extracellular concentrations of DA within the NAc. Cocaine extends DA transmission by blocking the DA transporter present on VTA dopaminergic neurons, and this mechanism is required for behavioral and physiological responses to cocaine^5^. Drug-induced elevations in DA promote transcriptional and epigenetic reorganization that ultimately gives rise to the cellular and behavioral adaptations that drive addiction-linked behaviors. Within the NAc, DA engages the type-1 DA receptor (DRD1, encoded by the *Drd1* gene) that is present on a subset of medium spiny neurons (MSNs), called D1-MSNs. DRD1 activation stimulates signal transduction cascades that results in CREB phosphorylation and activation of immediate early gene (IEG) expression programs. These IEG programs are comprised largely of activity-dependent transcription factors such as *Fos*, *FosB*, and *Nr4a1*^6–8^, each of which are critical mediators for cellular and behavioral adaptations to drugs of abuse^9–11^.

The NAc is a heterogeneous brain region consisting of many transcriptionally defined cell types. While a significant amount of research has focused on the role of DA receptor-expressing MSNs in adaptations to drugs of abuse, recent reports have also demonstrated significant roles for other cell types. For example, optogenetic inhibition of NAc somatostatin expressing interneurons (Sst-interneurons) attenuates cocaine-induced conditioned place preference^12^, suggesting that these cells are important for behavioral adaptations to drugs of abuse. Similarly, photoinhibition of cholinergic interneurons (Chat-interneurons) also blocks cocaine-induced conditioned place preference^13^. Interestingly, recent research has also demonstrated an important role for astrocytes in mediating cocaine-induced synaptic and behavioral adaptations^14,15^. These results demonstrate that diverse cell types within the NAc contribute to the molecular, cellular, and behavioral effects of cocaine.

Despite established cell type heterogeneity in the NAc, many studies rely on transgenic animal models that allow for the investigation of these changes only in individual cell populations. Unlike many techniques focused on a defined class of cells, single nucleus RNA-sequencing (snRNA-seq) provides unprecedented resolution that allows for the investigation of thousands of genes in thousands of cells with a single assay. This resolution not only permits detection of cocaine-induced transcriptional changes in many cell types, but also allows for better understanding of the molecular heterogeneity within the NAc. Recently, snRNA-seq approaches have been used to generate cellular atlases of the mouse^16^, non-human primate^17^, and human^18^ NAc. However, snRNA-seq has only been used to define early gene expression changes to acute cocaine, while the transcriptional effects of repeated cocaine in various NAc cell types is less well studied. Here, we used snRNA-seq to transcriptionally profile 39,254 nuclei from male and female Sprague-Dawley rats. Rats were further subdivided by treatment (cocaine or saline) and exposure paradigm (acute or repeated). This study confirms both the presence of many previously defined cell types within the rat NAc, and reveals that D1-MSNs exhibit the most robust transcriptional response to cocaine across all cell populations. Additionally, snRNA-seq of the rat NAc allowed for the investigation of cluster-specific expression of different neurotransmitter receptor systems and identified novel marker genes for two populations of transcriptionally distinct D1-MSNs.

## RESULTS

### Identification of transcriptionally distinct cell types within the rat NAc

To begin understanding cell-type heterogeneity within the rat NAc, we first integrated 15,655 nuclei from a previously published dataset^6^ consisting of male and female adult Sprague-Dawley rats that received a single (acute) injection of cocaine with 23,599 nuclei from a new dataset following seven daily (repeated) injections of cocaine (**Fig. 1a, Supp. Fig. 1)**. Following integration and dimensionality reduction, we identified 11 transcriptionally distinct neuronal populations and 5 glial populations that were not specific to a single dataset and did not differ by sex (**Fig. 1b, c, Supp. Fig. 2, 3**). Furthermore, calculation of Local Inverse Simpson’s Index (LISI) demonstrated that all clusters were well mixed, indicating successful integration (**Supp. Fig. 2**). A significant portion of neurons within the NAc are GABAergic MSNs that express high levels of *Gad1*, *Ppp1r1b*, *Foxp2*, and *Bcl11b* (**Fig. 1d, e**). A large majority of MSNs can be further classified by the expression of *Drd1*, the gene encoding the type-1 dopamine receptor, or *Drd2*, the gene encoding the type-2 dopamine receptor (**Fig. 1d**). While we have previously reported the presence of two transcriptionally distinct *Drd2*-expressing MSN populations (Drd2-MSN-1 and Drd2-MSN-2)^6^, this analysis also identified two transcriptionally distinct *Drd1*-expressing MSN populations as well (Drd1-MSN-1 and Drd1-MSN-2; **Fig. 1b,d**). In addition to Drd1- and Drd2-MSNs, this analysis also identified previously reported *Drd3*- and *Grm8*-expressing MSNs (Drd3-MSN and Grm8-MSN), which have also been identified within the striatum of non-human primates^17^. While most of the GABAergic neurons in the NAc are MSNs, distinct populations of GABAergic interneurons are also known to play critical yet distinct roles in striatal function^19^. While we have previously characterized parvalbumin and somatostatin interneurons within the NAc^6^, this analysis also identified a cholinergic interneuron population marked by the expression of *Chat* (Chat-Interneuron). Previous research has demonstrated that Chat-Interneurons are critical regulators of drug-induced behavioral adaptations^13^, and this dataset now provides comprehensive transcriptional profiling of these cells within the rat NAc. In addition to transcriptionally distinct neuronal populations, we also identified 5 glial clusters that are almost entirely devoid of *Gad1* and *Syt1* mRNA, and expressed canonical markers of their respective populations (**Fig. 1d,e**). For example, Astrocytes exhibit high expression of both *Aqp4* and *Gja1*, while Oligodendrocytes exhibit high expression of *Mbp*, the gene encoding myelin basic protein (**Fig. 1d, e**).

**Figure 1.**
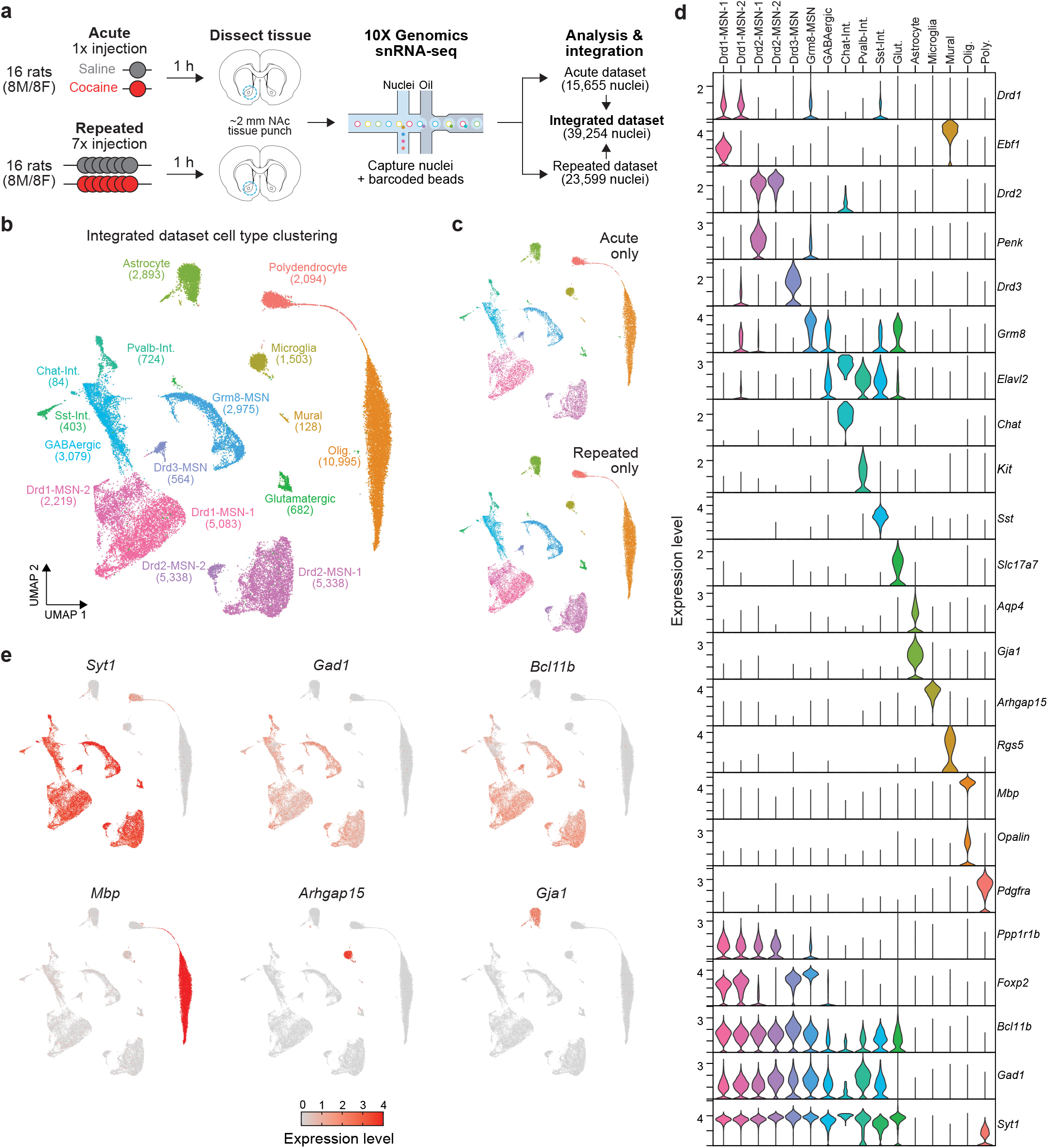
Experimental design and NAc cell cluster identification. **a**, Treatment paradigms and sample sizes for acute or repeated cocaine experiments. Rats (N=16/protocol, 8M/8F) received intraperitoneal injections of saline or cocaine (20mg/kg), and brain tissue was harvested 1 h after injections. NAc tissue was used for microfluidics-based nuclei capture, barcoding, and snRNA-seq on the 10X Genomics platform. The acute cocaine dataset was previously published in Savell, Tuscher, Zipperly, Duke, Phillips, et al., 2020 *Science Advances*. Repeated and acute cocaine datasets were integrated for further analysis. **b**, UMAP displaying 16 transcriptionally distinct cell types identified in the NAc from the integrated dataset (n=39,254 nuclei). **c**, UMAPs showing distribution of cells from acute and repeated cocaine experiments across all cell types. **d**, Violin plots displaying expression levels of marker genes of each cell type. **e**, Feature plots showing distribution of expression values across all cell types for marker genes for all neurons (*Syt1*), GABAergic cells (*Gad1*), medium spiny neurons (*Bcl11b*), oligodendrocytes (*Mbp*), microglia (*Arhgap15*), and astrocytes (*Gja1*).

### Computational investigation of transcriptionally distinct populations of Drd1-expressing MSNs

The integration of over 39,000 nuclei from rats that underwent different cocaine exposure paradigms identified two populations of *Drd1*-expressing MSNs (Drd1-MSN-1 and Drd1-MSN-2; **Fig. 2a).** To begin understanding the distinct gene expression signatures of these Drd1-MSN populations, we conducted a differential expression analysis using pseudobulked gene expression matrices. This analysis identified 506 genes enriched in the Drd1-MSN-1 cluster and 665 genes enriched in the Drd1-MSN-2 cluster. Interestingly, *Ebf1* is the most highly enriched differentially expressed gene in the Drd1-MSN-1 population (**Fig. 2b**). *Ebf1* is a known marker of D1-MSNs and is implicated in MSN early development^20^. *Calb1*, the gene encoding the calcium binding protein Calbindin 1, is another gene that is enriched in the Drd1-MSN-1 population (**Fig. 2b**). *Calb1* is a spatial marker within the NAc and is highly enriched in the NAc core^21^. Included in the top 10 differentially expressed genes in the Drd1-MSN-2 population was *Htr4*, the gene that codes for the serotonin receptor 4 (**Fig. 2b**). HTR4 autoradiography suggests that this receptor is located in the ventral portion of the NAc shell^22^. The differential enrichment of *Htr4* and *Calb1* suggests that the two populations of *Drd1*-expressing MSNs identified by snRNA-seq are both transcriptionally and spatially distinct.

**Figure 2.**
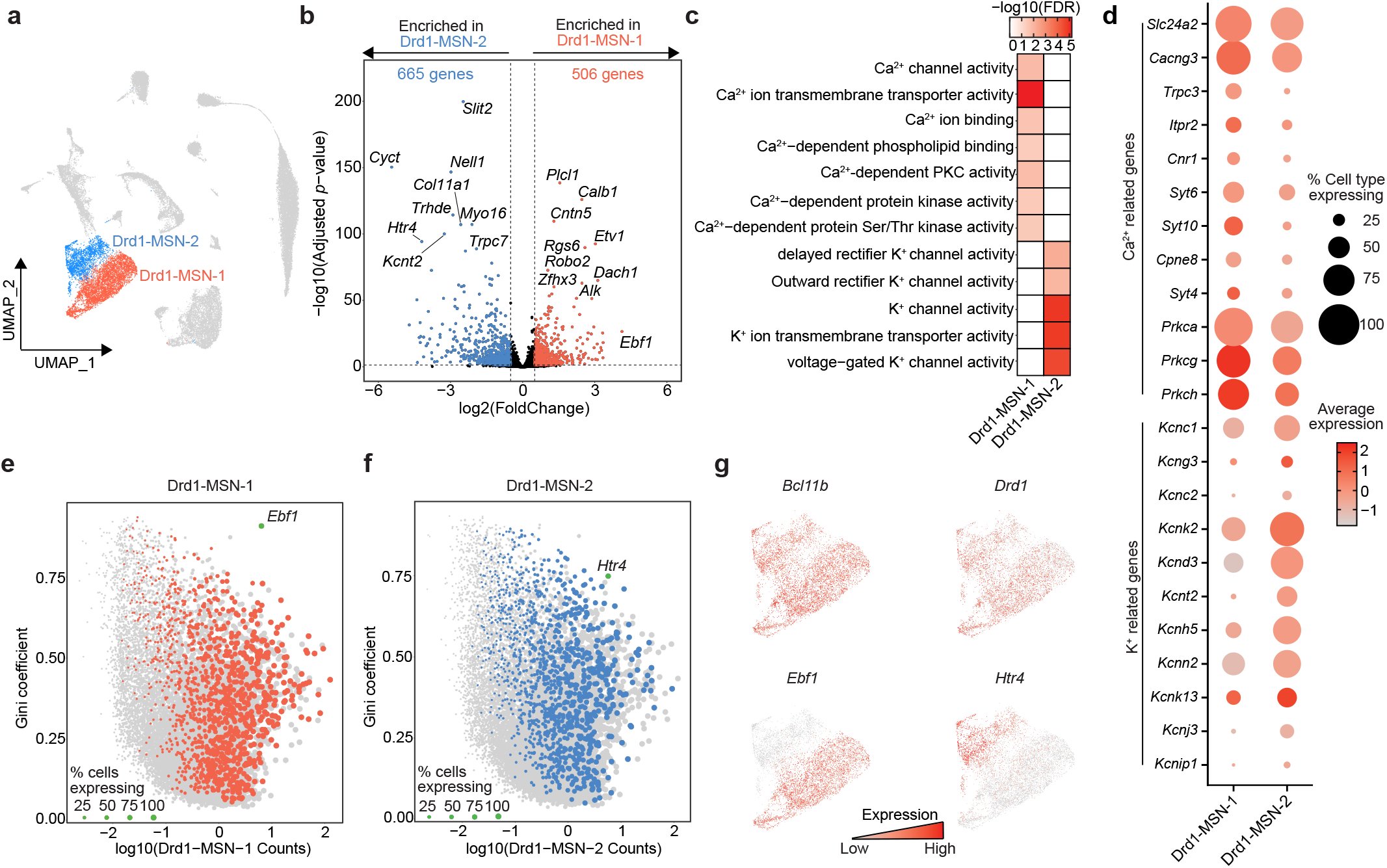
Identification of genes and pathways enriched in transcriptionally distinct populations of Drd1-expressing MSNs. **a**, UMAP highlighting the two identified populations of Drd1-expressing MSNs. **b**, Results of pseudobulk analysis directly comparing cells within the Drd1-MSN-1 population to cells within the Drd1-MSN-2 population. Enriched genes are those with an adjusted *p*-value < 0.05 and a |log2(Fold Change)| > 0.5. 506 genes were found to be enriched in the Drd1-MSN-1 population and are highlighted in red with a positive log2(Fold Change). 665 genes were found to be enriched in the Drd1-MSN-2 population and are highlighted in blue with a negative log2(Fold Change). **c**, Molecular function gene ontology analysis highlights enrichment of calcium-dependent processes in Drd1-MSN-1 population, while potassium-dependent processes are enriched in the Drd1-MSN-2 population. **d**, Dot plot showing expression levels and percent of cells within the clusters expressing genes involved in calcium- or potassium-dependent processes. **e-f**, Scatter plots highlighting differentially expressed genes with high counts and high Gini coefficients. This analysis identified *Ebf1* and *Htr4* as potential markers of Drd1-MSN-1 and Drd1-MSN-2, respectively. **g**, Feature plots showing enrichment of *Bcl11b* (MSN marker), *Drd1* (Drd1-MSN marker), *Ebf1* (Drd1-MSN-1 marker), and *Htr4* (Drd1-MSN-2 marker).

Next, we set out to understand the specific gene networks and pathways that may be enriched in these two D1-MSN populations. Molecular function pathway analysis revealed an enrichment of genes involved in calcium transport and signaling in the Drd1-MSN-1 population, but enrichment of genes involved in potassium transport in the Drd1-MSN-2 population (**Fig. 2c**). For example, *Trpc3*, a gene encoding a calcium permeable channel, and *Cacng3*, a gene encoding an L-type calcium channel, are both highly enriched in the Drd1-MSN-1 population (**Fig. 2d**). Conversely, several genes encoding potassium channels, such as *Kcnc1*, *Kcnc2*, and *Kcng3*, are enriched in the Drd1-MSN-2 population, further highlighting the enrichment of genes involved in potassium transport in this population. While these genes are differentially enriched, both calcium- and potassium-related genes are expressed to some extent in both clusters (**Fig. 2d**).

Functional investigation of transcriptionally distinct populations of neurons in the mammalian brain requires the use of cell type-specific promoters. To facilitate the future investigation of the two populations of transcriptionally distinct *Drd1*-expressing MSNs, we next identified potential marker genes that are both differentially and highly expressed within a population. This analysis identified *Ebf1* as a potential marker gene for the Drd1-MSN-1 population (**Fig. 2e**) and *Htr4* as a potential marker gene for the Drd1-MSN-2 population (**Fig. 2f**). An additional characteristic of marker genes is that they are expressed in a large proportion of cells within the cluster. Plotting the expression of these genes on a UMAP shows differential enrichment of *Ebf1* and *Htr4* between the *Drd1*-expressing MSN populations (**Fig. 2g**). However, *Ebf1* is also expressed in endothelial cells, while *Htr4* is also expressed in Drd2- and Drd3-MSNs. Thus, these genes may act as marker genes in combination with *Drd1* but cannot act as marker genes alone.

### In-situ validation of transcriptionally distinct Drd1-expressing MSNs

To validate snRNA-seq findings demonstrating that *Htr4* and *Ebf1* are in largely distinct populations of *Drd1*-expressing MSNs in the NAc, we used RNAscope^23^, a previously established fluorescence *in situ* hybridization (FISH) protocol. To enable visualization of distinct puncta representing cells expressing a combination of *Drd1*, *Htr4*, and *Ebf1*, we performed multiplexed analysis with probe sets specific to each transcript (**Fig. 3a-c**). Consistent with our computational analysis, we identify unique subpopulations of *Drd1*+ cells that are either *Ebf1*+ or *Htr4*+, with significantly fewer cells that are positive for both *Ebf1* and *Htr4* (**Fig. 3b-c**). Furthermore, our *in situ* experiments revealed unique spatial distributions for these subpopulations with a significantly higher proportion of *Drd1*+/*Htr4*+ cells in the NAc shell and *Drd1*+/*Ebf1*+ cells in the NAc core (**Fig. 3d-e**). There was no significant spatial distribution difference in the proportion of *Drd1+/Ebf1+/Htr4+* cells. Overall, these results are consistent with snRNA-seq analysis, and confirm largely non-overlapping expression of *Ebf1* and *Htr4* within *Drd1*+ cells (**Fig. 3f**).

**Figure 3.**
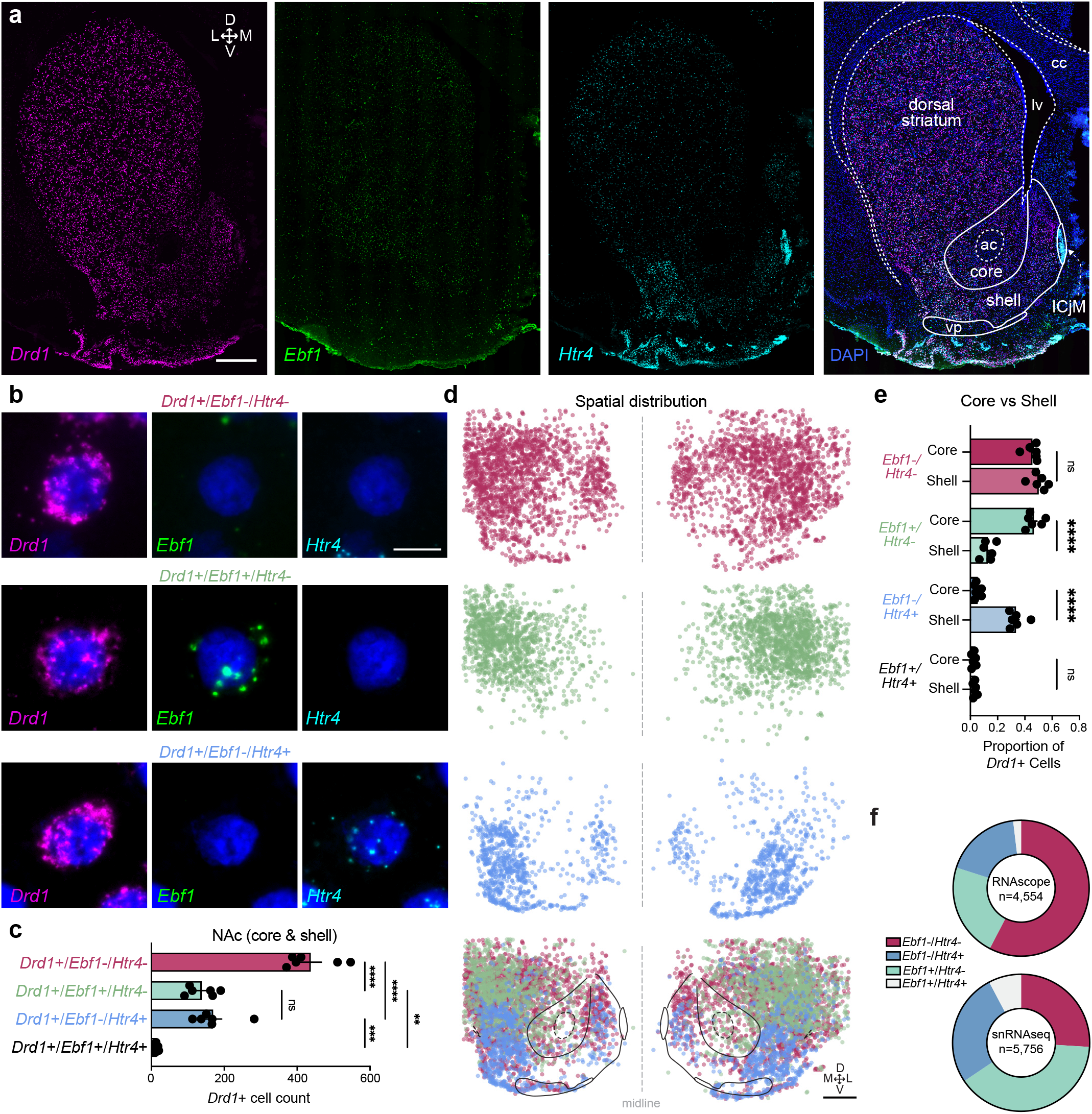
*In situ* validation of transcriptionally distinct populations of *Drd1*-expressing MSNs. **a**, Representative 20x stitched image following RNAscope in situ hybridization with probes for *Drd1*, *Ebf1*, *Htr4*, and DAPI. Scale bar: 1mm. ac: anterior commissure; cc: corpus callosum; lv: lateral ventricle; vp: ventral pallidum. ICjM: Islands of Calleja Major. Scale bar: 1mm. **b**, 100x images showing representative *Drd1*+/*Ebf1*-/*Htr4*-, *Drd1*+/*Ebf1*+/*Htr4*-, *Drd1*+/*Ebf1*-/*Htr4*+, and *Drd1*+/*Ebf1*+/*Htr4*+ cells, with *Drd1*, *Ebf1*, *Htr4*, and merged channels. Scale bar: 10μm. **c**, Quantification and comparison of *Drd1*+ subpopulations in the NAc. One-way ANOVA test with Tukey’s multiple comparison test. **d**, Location of cells expressing selected genes and anatomical overlay. **e**, Quantification and comparison of *Drd1*+ subpopulations in the NAc core and shell. One-way ANOVA test with Tukey’s multiple comparison test. **f**, Proportion of *Drd1*+ cell types in the NAc from RNAscope (n=6 animals) and snRNAseq datasets. \**p*<0.05, \*\**p*<0.01, \*\*\**p*<0.001, \*\*\*\**p*<0.0001, ns: not significant.

### Cocaine experience differentially alters the transcriptome of specific cell types within the NAc

We previously demonstrated that a single acute cocaine exposure differentially alters the transcriptome of molecularly defined cell types within the NAc^6^. To begin understanding how repeated cocaine exposure alters the transcriptome of distinct NAc populations, we injected male and female Sprague-Dawley rats with 20mg/kg cocaine for 7 consecutive days (**Fig. 1a, Supp. Fig. 1**). Before directly comparing acute and repeated exposures, we first investigated cell type-specific transcriptional responses to cocaine exposure, regardless of type or duration. To do this, we collapsed both datasets and used a pseudobulk approach to identify cocaine-dependent, cell type-specific differentially expressed genes (DEGs; **Supp. Table 2**). This approach increased the number of replicates used for statistical comparisons and circumvented previously identified issues with deprecated differential expression methods previously used for snRNA-seq^24^. Strikingly, the Drd1-MSN-1 population exhibited the most robust transcriptional response to cocaine, with 236 total DEGs following cocaine experience (184 upregulated, 52 downregulated; **Fig. 4a, b**). These upregulated DEGs include immediate early genes (IEGs) such as *Nr4a3*, *Tiparp*, and *Fosl2* (**Fig. 4b**). Recent evidence has demonstrated that astrocytes within the NAc also play a key role in the neuronal and behavioral response to dopamine and drugs of abuse^14,15,25^. Disruption of glutamatergic transport, release, and signaling is implicated in substance use disorders^26,27^, and astrocytes are heavily involved in the regulation of this process across the brain^28^. Interestingly, *Gria1*, an ionotropic AMPA receptor, is significantly decreased in Astrocytes following cocaine exposure (**Fig. 4c**). Unlike the Drd1-MSN-1 population, Astrocytes exhibited more genes with cocaine-dependent decreases in expression (**Fig. 4c,g**). While most DEGs in the Astrocyte population are downregulated with cocaine, *Pde10a*, the gene encoding the cAMP-regulated phosphodiesterase 10A, is upregulated by cocaine in both the Astrocyte and Sst-Interneuron cluster (**Fig. 4c,d**).

**Figure 4.**
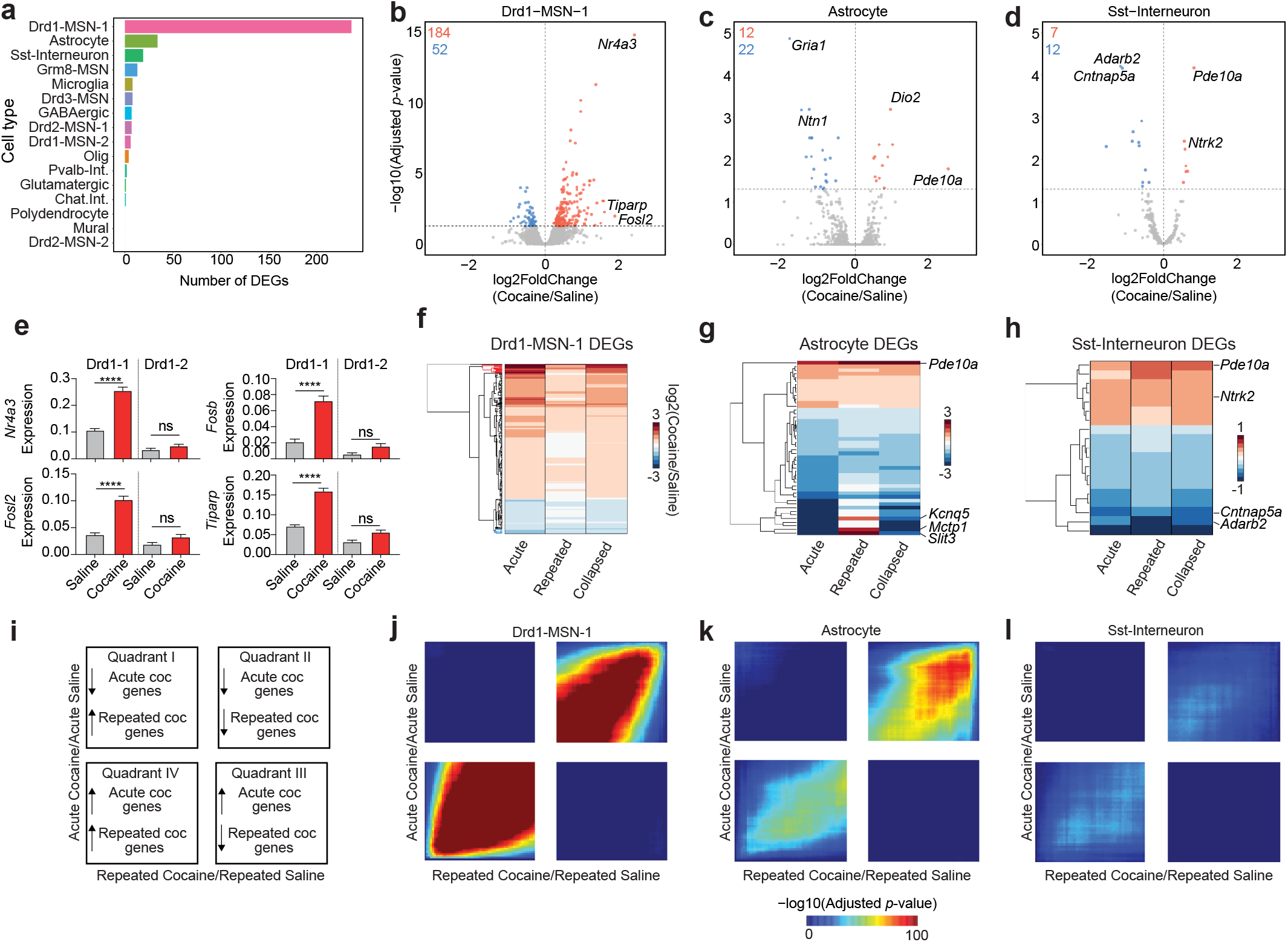
Cocaine-dependent transcriptional changes identified using pseudobulk differential expression analysis. **a**, Number of cocaine dependent DEGs in each identified cell type. **b-d**, Volcano plots for Drd1-MSN-1, Astrocyte, and Sst-Interneuron populations. **e**, Cocaine dependent changes for 4 different immediate early genes in the Drd1-MSN-1 and Drd1-MSN-2 populations (One-way ANOVA test with Tukey’s multiple comparison test:\*\*\*\**p*<0.0001, ns: not significant). **f-h**, Heatmap showing log2(Fold Change) of all cocaine DEGs in Drd1-MSN-1, Astrocyte, and Sst-interneuron populations, regardless of length of cocaine exposure. **i**, Interpretation key for rank-rank hypergeometric overlap (RRHO) heatmaps. Quadrants I and III represent discordant gene expression changes, while quadrants II and IV represent concordant gene expression changes. **j-l**, RRHO heatmaps for Drd1-MSN-1, Astrocyte, and Sst-Interneuron.

Previous research has demonstrated that *Drd1*-expressing MSNs exhibit robust transcriptional changes in response to cocaine^6,8,9,29^. More specifically, snRNA-seq from the rat NAc following acute cocaine exposure demonstrated that D1-MSNs have the most robust transcriptional response to cocaine when compared to other transcriptionally distinct NAc cell types^6^. In this dataset, only one of the *Drd1*-expressing MSN populations (Drd1-MSN-1) had a robust transcriptional response to cocaine. For example, *Nr4a3*, *Fosb*, *Fosl2*, and *Tiparp* are all significantly increased following cocaine exposure only in the Drd1-MSN-1 population (**Fig. 4e**).

Next, we set out to investigate the relationship of cocaine dependent DEGs across cocaine exposure paradigms. To do this, we first conducted DEG analyses comparing cocaine and saline samples with exposure type ignored (i.e., all cocaine samples vs all saline samples), acute exposure only, and repeated exposure samples only. The log2 (Fold Change) values for all DEGs were then concatenated and subject to hierarchical clustering. Overall, this analysis identified that many of the genes altered by acute cocaine are also altered by repeated cocaine in the same direction (**Fig. 4f-h**). However, for the Drd1-MSN-1 population some genes altered by acute cocaine are not altered by repeated cocaine. Furthermore, while the overall pattern shows concordant gene expression patterns, some cocaine-regulated genes within the Astrocyte cluster demonstrate opposite changes with an acute or repeated exposure. To estimate the degree of concordance between gene expression programs altered by acute and repeated cocaine exposure in a threshold-free manner, we next used the rank-rank hypergeometric overlap (RRHO) algorithm^30^. To complete this analysis, we ranked gene lists by the log2 (Fold Change), which was calculated by dividing the log-normalized expression values of cocaine cells by the log-normalized expression values of saline cells. This analysis identified that, on a cell type-specific basis, acute and repeated cocaine gene expression signatures are primarily concordant. For example, the acute and repeated cocaine changes for the Drd1-MSN-1, Astrocyte, and Sst-Interneuron populations are significantly concordant for both upregulated and downregulated genes (**Fig. 4i-l**). However, the strength of this overlap and concordance is substantially different for these populations.

### Conservation of MSN subtypes across mammalian species

Recently, transcriptional atlases of the non-human primate (NHP) striatum^17^ and human NAc^18^ have demonstrated similar gene expression profiles of specific marker genes that we identify within the rat NAc. To quantify this overlap and perform cross-species comparisons, we focused on conservation of MSNs given that both NHP and human datasets identified and validated the presence of several transcriptionally distinct MSN subtypes. Direct comparison of our rat snRNA-seq dataset to monkey or human datasets revealed potential conservation of many NHP and human cell types (**Fig. 5a-c**). For example, the rat Drd1-MSN-1 population is correlated with the human “D1. MSN_A” and NHP “D1.Matrix” populations, and the rat Drd2-MSN-1 population is correlated with the human “D2.MSN_A” and the NHP “D2.Matrix” population (**Fig. 5a-d**).

**Figure 5.**
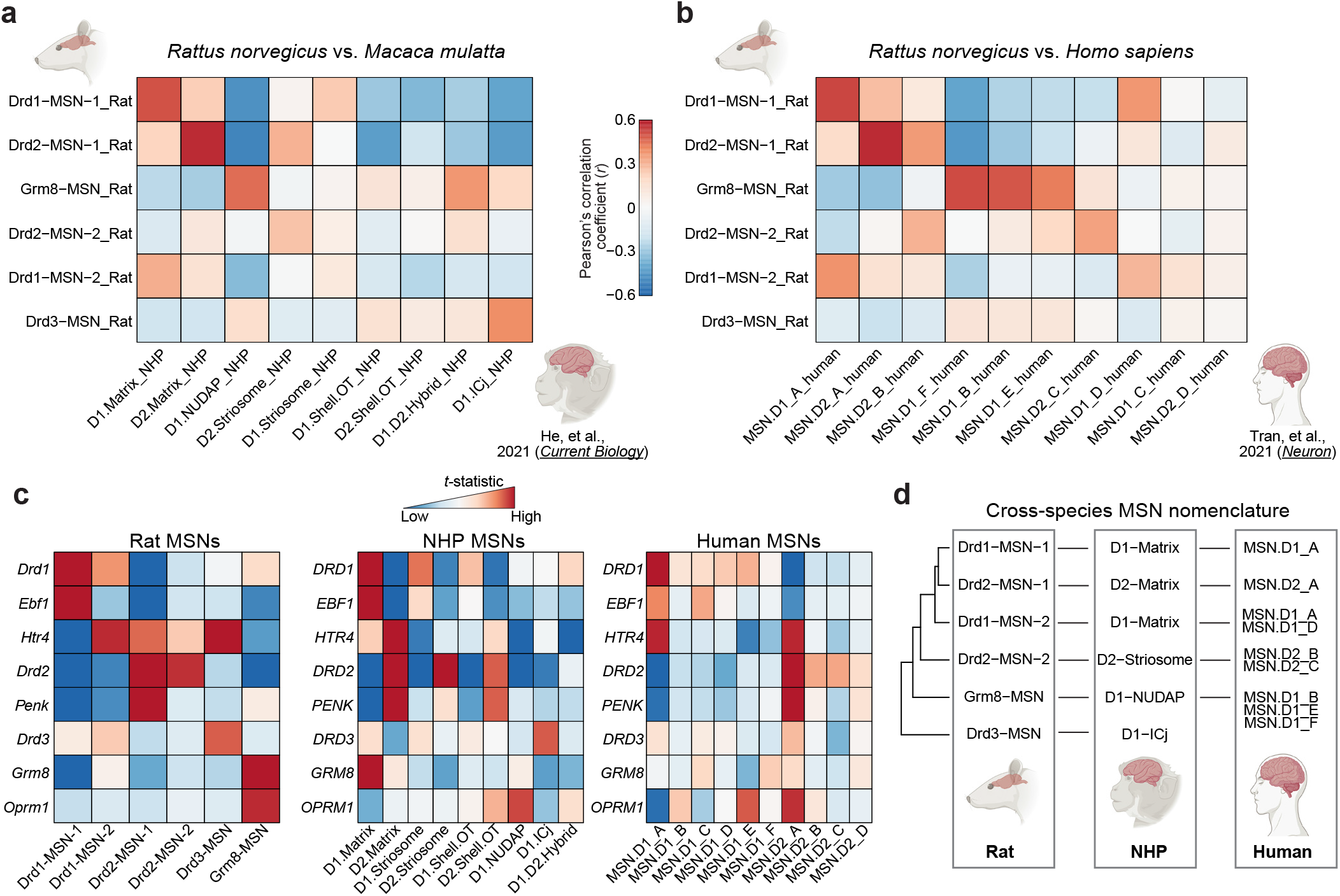
Identification of conserved neuronal cell types across higher order mammals. **a-b**, Heatmap colored by the Pearson’s correlation coefficient between rat, non-human primate, and human cell types. Correlation coefficients were calculated using t-statistic for rat MSN DEGs and their orthologs. **c**, Heatmaps colored by *t*-statistic for specific MSN marker genes in rat, non-human primate, and human. **d**, Estimated conservation of NAc neuronal cell types across rat, non-human primate, and human.

Transcriptionally conserved cell types should exhibit similar expression and enrichment of marker genes. For example, Drd2-MSNs are classically characterized by the enrichment of *Drd2* and *Penk*, while Drd1-MSNs are characterized by the enrichment of *Drd1* and *Ebf1*. Across species, transcriptionally distinct populations of Drd2-MSNs are similarly enriched for *Drd2*, but differ in their enrichment of *Penk* (**Fig. 5c, Supp. Fig. 4a-c)**. A similar pattern is observed in Drd1-MSNs across species, as *Drd1* is highly enriched in Drd1-MSNs, but *Ebf1* enrichment differs (**Fig. 5c, Supp. Fig. 4a-c).** While the differential expression of *Htr4* or *Ebf1* distinguishes Drd1-MSNs in the rat NAc, this pattern is not observed in higher order mammals as *Htr4* is ubiquitously expressed across cell types (**Supp. Fig. 4a-c**). Furthermore, *Htr4* is enriched in the “MSN.D1_A” human population (**Fig. 5c**) along with *Ebf1*. Thus, *Htr4* and *Ebf1* may not be suitable markers for distinguishing between transcriptionally distinct populations of Drd1-MSNs in higher order mammals.

## DISCUSSION

Within the past few decades, a major focus of drug addiction research has been on the transcriptional and epigenetic changes occurring in the NAc following drug exposure. While these studies have been integral in our understanding of how drug experience shapes the brain, they have primarily used techniques that ignore cell-type heterogeneity. Furthermore, although the NAc is fundamental in reward processing and the rat is a primary model organism used for reward-related research, few studies have comprehensively profiled the rat NAc at the transcriptional level. snRNA-seq provides the unique opportunity to study both the cell type-specific transcriptional responses to drugs of abuse, as well as the cell-type heterogeneity that exists within the NAc. In the present study, we report the integration of a previously published NAc snRNA-seq dataset collected following acute cocaine exposure^6^ with a new dataset collected following repeated cocaine exposure. The resulting dataset contains 39,254 nuclei across 16 transcriptionally distinct populations (11 neuronal and 5 non-neuronal), which more than doubles the size of our previously published work. The addition of both biological and technical replicates, along with the addition of over 24,000 nuclei, allowed us to generate a comprehensive transcriptional atlas of the rat NAc and investigate cell type-specific transcriptional responses to cocaine.

We were also interested in understanding the distribution of expression of genes encoding different receptor systems, which provides additional information regarding which cell types may respond to diverse classes of drugs. For example, there is significant renewed interest in the action of psychedelic drugs, many of which act on serotonin receptors. *Htr2a*, the gene encoding serotonin receptor type 2A and the expected target of many psychedelics, is expressed in a small proportion of *Drd1*- and *Drd2*-expressing MSNs, as well as Sst-interneurons (**Supp. Fig. 5a**). Thus, these populations of cells will most likely respond to classic psychedelics. There is also considerable interest in the distribution of expression of opioid receptors in the NAc as opioid-associated overdose deaths continue to increase^2,3^. Previous research has demonstrated that 25% of cells containing the delta opioid receptor (*Oprd1*) in the NAc also contained Met^5^-Enkaphalin^31^, a peptide encoded by the gene *Penk. Penk* is a marker gene for *Drd2*-expressing MSNs, suggesting that the delta opioid receptor is expressed in this population of neurons. Our data confirms this observation, as *Oprd1* is highly expressed in the Drd2-MSN-1 population (**Supp. Fig. 5a-c**).

A major focus of this study was to characterize cell type-specific gene programs induced by cocaine regardless of exposure length. Independent of exposure type, the largest transcriptional response to cocaine experience occurred in the Drd1-MSN-1 population (**Fig. 4a, b**). These changes included IEGs such as *Fosl2*. IEGs and other cocaine-regulated genes in the Drd1-MSN-1 population are induced by both acute and repeated cocaine, but to a lesser extent with repeated cocaine. This data is in agreement with previous research that demonstrated induction of FOSB protein by repeated cocaine was weaker as compared to acute cocaine^29^. The second population of *Drd1*-expressing MSNs within the NAc, Drd1-MSN-2, did not have a robust transcriptional response to cocaine experience. A multitude of previous research has demonstrated that *Drd1*-expressing cells within the NAc exhibit transcriptional and epigenetic responses to drugs of abuse^6,8,9,29,32^. Thus, the identification of a population of *Drd1*-expressing MSNs in the NAc with a significantly attenuated transcriptional response to cocaine is intriguing because it highlights a previously underappreciated level of molecular heterogeneity within this population of cells. Notably, further analysis of population-specific DEGs also revealed an enrichment of calcium transport-related terms for Drd1-MSN-1 and potassium transport-related terms for Drd1-MSN-2. The differential enrichment for calcium-related terms and genes may explain the transcriptional response to cocaine in the Drd1-MSN-1 population, as many IEGs rely on calcium-dependent signaling cascades for induction.

Our dataset also provided the opportunity to identify marker genes for these two populations. We identified *Ebf1* as a marker gene for the Drd1-MSN-1 population and *Htr4* as a marker gene for the Drd1-MSN-2 population. *In situ* experiments validated the presence of two distinct populations of *Drd1*-expressing MSNs based on the presence of *Htr4* and *Ebf1*. These *in situ* experiments also revealed a spatial separation of these MSNs, as *Drd1*+/*Ebf1*+ cells were primarily found in the NAc core and *Drd1*+/*Htr4*+ cells were primarily found in the NAc shell. Transcriptional similarity based on spatial location may also explain why the Drd1-MSN-1 population is more closely related to the Drd2-MSN-1 population than the Drd1-MSN-2 population (**Supp. Fig. 4a**). In fact, analysis of the relationship between cell types within the NHP identifies that *Drd1*- and *Drd2*-expressing MSNs from the same striatal subregions are more closely related than *Drd1*-expressing MSNs found in distinct striatal compartments (**Supp. Fig. 4b**). Similarly, the MSN.D1_A and MSN.D2_A human populations are more closely related to each other than other *Drd1*- or *Drd2*-expressing MSNs, although no spatial information is available in that dataset (**Supp. Fig. 4c)**. Together, these results suggest that at least some component of transcriptional diversity in the NAc reflects the spatial organization of this structure.

Although sex differences in drug-induced behavioral adaptations have long been appreciated^33–36^, more recent research has also demonstrated sex-specific drug-induced molecular adaptations^37^. Our dataset includes nuclei from both male and female rats, and thus we examined sex-specific transcriptional responses to cocaine across cell populations within the NAc, again using the collapsed acute and repeated dataset to enable pseudobulk treatment comparisons between male and females. In general, the direction of cocaine-induced mRNA changes was largely concordant between males and females across most cell populations (**Supp. Fig. 6a-d**), however significance criteria were often only met for one sex (**Supp. Table 3**). Overall, the Drd1-MSN-1 population was the most transcriptionally responsive to cocaine for both sexes in terms of total number of genes altered. Consistent with the sex-combined identification of cocaine DEGs, immediate early gene *Nr4a3*, transcription elongation factor *Ell2*, and cytoskeletal regulator *Elmo2* were all significantly elevated in the Drd1-MSN-1 population in both sexes (**Supp. Fig. 6e**). Interestingly, sex-specific cocaine DEG comparisons revealed that while some genes, like *Nr4a3* and *Ell2*, display relatively consistent patterns of expression between males and females across populations (**Supp. Fig. 6e-f**), other genes like *Hs6st3* exhibit more dynamic patterns of expression (**Supp. Fig. 6g**).

In conclusion, we have generated a comprehensive transcriptional atlas of the rat NAc that also provides information regarding cocaine-dependent gene expression changes. The use of snRNA-seq provided the ability to understand cell types that potentially mediate cellular and behavioral responses to drugs of abuse. Analysis of this dataset resulted in the identification of transcriptionally distinct populations of *Drd1*-expressing MSNs that differ by their spatial location and transcriptional response to cocaine. This dataset will be a resource for future research into transcriptionally distinct populations of NAc cells and their responses to drugs of abuse.

## METHODS

### Behavioral testing

All experiments were performed in accordance with the University of Alabama at Birmingham Institutional Animal Care and Use Committee. Male or female adult rats were intraperitoneally injected with Cocaine hydrocholoride at a dose of 20mg/kg or saline. Locomotor testing was performed as previously described^6^. Animals undergoing acute cocaine exposure received a single dose, while those undergoing repeated cocaine exposure received one injection per day for 7 days.

### Single-nuclei dissociation

Nuclei dissociation for snRNA-seq was performed as previously described^6^. Nuclei from 4 animals of the same sex, treatment group, and exposure length were pooled for tissue preparation and loaded into a single GEM (gel beads in emulsion) well.

### snRNA-seq Analysis Methods

Libraries were constructed according to manufacturer’s instructions using the Chromium Single Cell 3’ Library Construction Kit (10x Genomics, catalog no. 1000092), which uses version 3 chemistry for gene expression. Nuclei (15,655) from adult rat NAc after acute cocaine and nuclei (23,599) from adult rat NAc after repeated cocaine were sequenced on the Illumina NextSeq 500 at the UAB Heflin Genomics core to a depth of ~39,000 reads per nuclei and ~22,000 reads per nuclei, respectively. All raw fastq files were aligned to the Ensembl mRatBn7.2 (Rn7) genome using CellRanger (v6.1.2) with the associated Ensembl gene transfer format (gtf) file (version 105). Cell Ranger filtered outputs were analyzed with Seurat v4.0.4^38^ using R v4.0.2. Nuclei containing <200 features and >5% of reads mapping to the mitochondrial genome were removed from further analysis. Molecular count data from each GEM (gel beads in emulsion) well were then log-normalized with a scaling factor of 10,000. Following normalization, and before dimensionality reduction, all GEM wells were then integrated with FindIntegrationAnchors() and IntegrateData using 17 principal components (PCs) and a resolution value of 0.1 for both the acute and repeated datasets. Local Inverse Simpson’s Index was calculated to ensure successful integration of all 8 GEM wells using lisi v1.0 with default parameters. Recent research has demonstrated that ambient RNA can be captured with single cells or nuclei with droplet-based single-cell/single-nucleus technologies. To avoid interpreting any signal associated with ambient RNA, we used SoupX v1.5.2^39^ to remove any signal associated with ambient RNA. Droplet based single-cell/single-nucleus technologies are also susceptible to the capture of “doublets”, or two cells/nuclei being captured within a single droplet. Homotypic doublets, or doublets containing two cells of the same cell-type, are difficult to identify and remove. However, heterotypic doublets, or doublets containing two cells of different cell types, can be identified as cells that have a combination of cell type-specific transcriptional signatures. Heterotypic doublets were not identified in our acute dataset. However, heterotypic doublets were identified and removed from the repeated dataset using DoubletFinder v2.0.3^40^. For differential expression testing, gene counts were first summed across GEM wells within each cell type to create a pseudobulked gene expression matrix. Any genes with counts <5 were removed from further analysis. Differential expression testing was then completed using pseudobulked gene expression matrices with DESeq2 v1.28.1^41^ using the Likelihood Ratio Test (LRT). To identify differentially expressed genes across genetic sex and cell types, dataset and treatment were used as covariates. To identify differentially expressed genes across treatment conditions, dataset was used as a covariate. Gini coefficients were calculated using the gini() function that is available in the reldist v1.6-6 package^42,43^. Rank-rank hypergeometric overlap (RRHO) was calculated using the log2 (Fold Change) of all genes regardless of adjusted p-value. RRHO heatmaps were generated using RRHO2 v1.0^30,44^. Molecular function gene ontology terms were generated with DEGs expressed in >10% of the cluster using g:GOSt made available by gProfiler^45^. For this tool, the statistical domain scope set to “Only annotated genes” and significance threshold set to “Benjamini-Hochberg” FDR. All R code for this analysis is available at https://github.com/Jeremy-Day-Lab/Phillips_2023_NAc.

### Calculation of t-statistic for conservation analysis

To correlate gene expression signatures across mammalian species, DEGs for rat cell types were first calculated using a pseudobulk approach. DEGs were then concatenated and t-statistics for every gene in every MSN subtype were calculated. The *t*-statistic was calculated with the following formula:

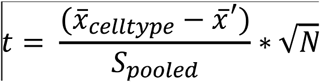

where 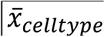 is the mean expression value of the gene in the cell type of interest, 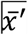 is the mean expression value of all other cells, *n* is the number of nuclei in the cell type of interest, and 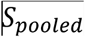 is the pooled standard deviation according to:

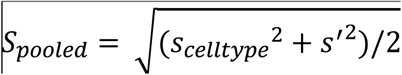

Following calculation of t-statistics, the Pearson correlation coefficient was calculated using the cor() function in R. All R code for this analysis is available at https://github.com/Jeremy-Day-Lab/Phillips_2023_NAc.

### Tissue Preparation and RNAscope

Upon sacrifice, brains from 6 drug-naïve Sprague-Dawley rats (3F/3M) were extracted and immediately placed in isopentane at −40°C for 30 seconds. Tissue was stored at −80°C until day of sectioning. On day of sectioning, tissue was allowed to equilibrate at −20°C for 40 minutes and 10μm coronal sections containing the NAc (1.92mm to 1.44mm from bregma) were collected using Leica cryostat and mounted on room temperature slides. Sections were air dried on dry ice for 60 minutes, then stored at −80°C. Sections were prepared for RNAscope Multiplex Fluorescent v2 Assay according to manufacturer’s protocol (Advanced Cell Diagnostics, 323110)^23^. Probes included *Htr4*-C2 (544081-C2), and *Drd1a*-C3 (317031-C3) diluted 1:50 in *Ebf1*-C1 (891561-C1) and DAPI (320858). Associated Opal dyes were 690 (FP1497001KT) for *Ebf1*-C1, 570 (FP1488001KT) for *Htr4*-C2, and 520 (FP1487001KT) for *Drd1a*-C3 diluted 1:500 in TSA buffer. Sections were imaged using Keyence BZ-X800 microscope with OP-87767 filter cubes for GFP, Texas Red PE, DAPI, and Cy5 at 20x (PlanFluor NA 0.75) and 100x (PlanApo NA 1.45) magnifications. 20x images were stitched using the BZ-X800 Analyzer software.

### Image Analysis

Stitched images were analyzed using ImageJ2 V2.9.0/1.53t^46^. Briefly, NAc-specific nuclei ROIs were generated based on DAPI channel image for each section using StarDist2d Versatile (fluorescent nuclei) plugin^47^. DAPI ROIs were applied to the other channels containing signals associated with *Ebf1*, *Htr4*, and *Drd1a* probes. Signal intensity for each channel, normalized and rescaled from 0-100 for each animal and probe, was measured to identify *Drd1a*+, *Drd1a*+/*Ebf1*+, *Drd1a*+/*Htr4*+, and *Drd1a*+/*Htr4*+/*Ebf1*+ populations.

## DATA AVAILABILITY

All snRNA-seq datasets have been deposited on NCBI GEO for download (GSE137763, GSE222418). A publicly accessible and searchable online resource for these datasets is available at http://day-lab.org/resources.

## ACKNOWLEDGEMENTS

We thank all current and former Day Lab members for assistance and support. This work was supported by NIH grants DP1DA039650, R01MH114990, R01DA053743, and R01DA054714 and the McKnight Foundation Neurobiology of Brain Disorders Award (JJD). L.I. is supported by the Civitan International Research Center at UAB. We acknowledge support from the University of Alabama at Birmingham Biological Data Science Core (RRID:SCR_021766), the UAB Heflin Center for Genomic Sciences, and the UAB Flow Cytometry & Single Cell Core Facility. We would also like to thank Nathaniel J. Robinson for his input for calculating the *t*-statistic from gene expression data for cross-species comparisons.

## AUTHOR CONTRIBUTIONS

R.A.P., J.J.T., M.E.Z, C.G.D., and J.J.D. conceived of sn-RNA-seq experiments. R.A.P., J.J.T., M.E.Z., & C.G.D. performed snRNA-seq tissue preparation. R.A.P., J.J.T., E.W., L.I., and J.J.D. performed bioinformatics analysis. R.A.P. and J.J.D wrote the manuscript with assistance from J.J.T. and N.D.F. N.D.F. performed RNAscope experiments and imaging. J.J.D. supervised all work. All authors have approved the final version of the manuscript.

---

**Supplementary Figure 1.**
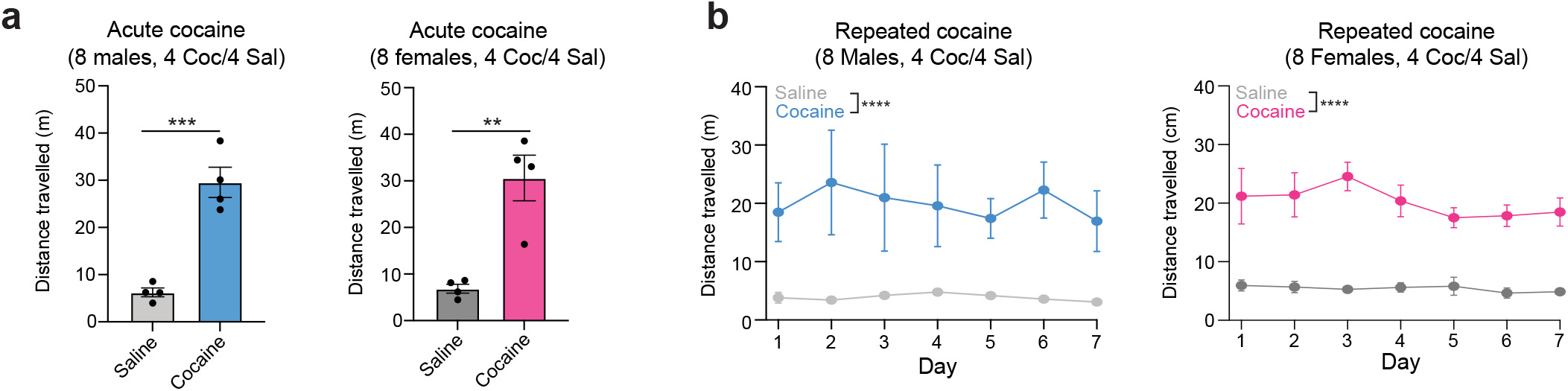
Cocaine-induced locomotor changes. **a**, Male and female Sprague-Dawley rats were treated with 20mg/kg for 1 hour and locomotor behavior was measured for 30 minutes. Cocaine-treated animals travelled signficantly more than saline-treated animals. Left, acute cocaine-induced locomotor changes in male rats (n=4/group, unpaired Student’s *t*-test: *t*(6)=6.974, \*\*\**p*<0.0005). Right, acute cocaine-induced locomotor changes in female rats (n=4/group, unpaired Student’s *t*-test: *t*(6)=4.769, \*\**p*<0.005) Data from Savell*, Tuscher*, Zipperly*, Duke*, Phillips* et al. 2020, *<u>Science Advances</u>*. **b**, Male and female Sprague-Dawley rats were treated with 20mg/kg once daily for 7 consecutive days. Locomotor behavior was measured for 30 minutes. Cocaine-treated animals travelled signficiantly more than saline-treated animals on each testing day. Left, repeated cocaine-induced locomotor changes in male rats across 7 days (n=4/group, two-way ANOVA for main effect of treatment F_(1,42)_ = 41.40, \*\*\*\**p*<0.0001). Right, repeated cocaine-induced locomotor changes in female rats across 7 days (n=4/groups, two-way ANOVA for main effect of treatment, F_(1,42)_ = 156.8, \*\*\*\**p*<0.0001).

**Supplementary Figure 2.**
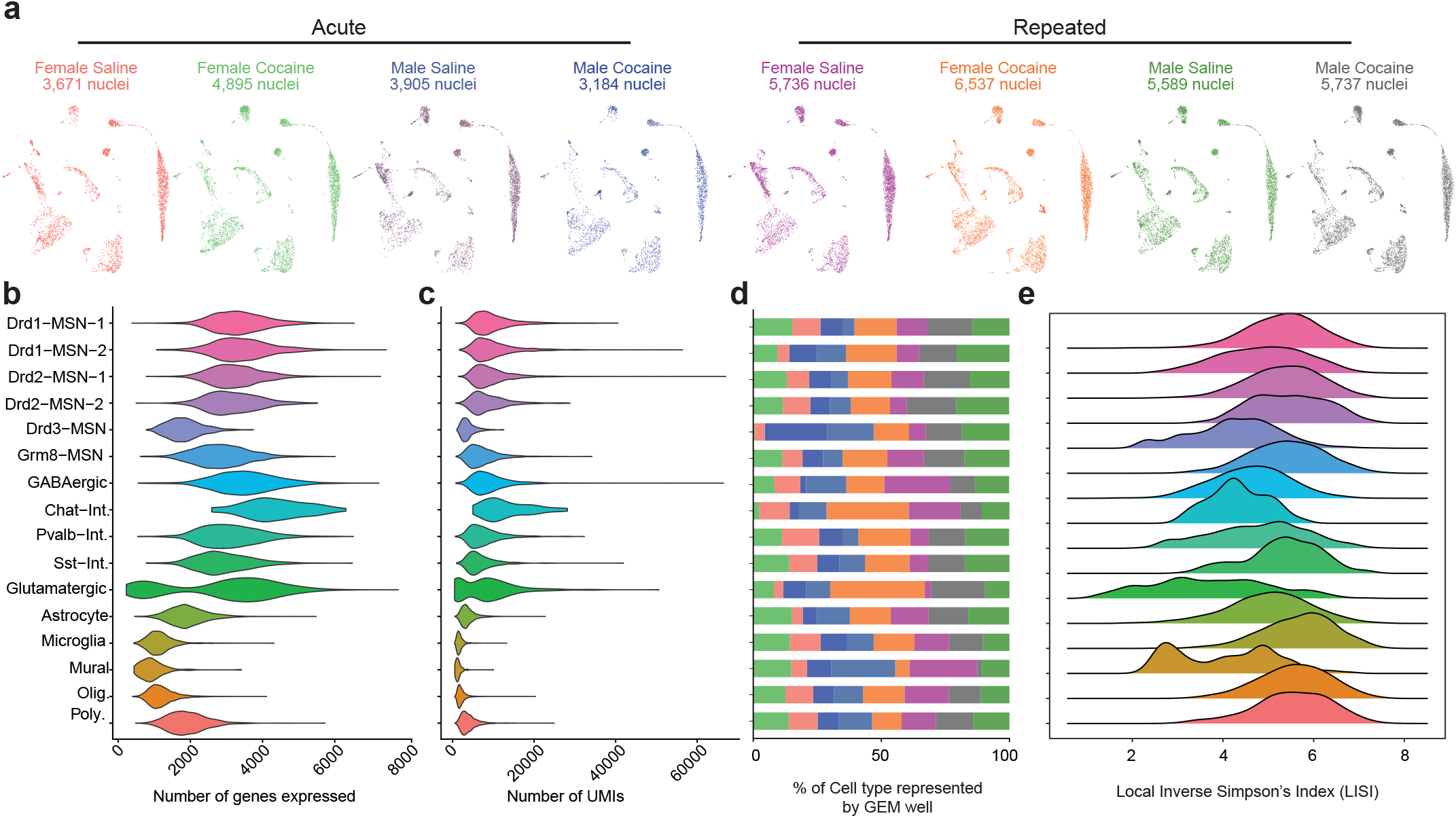
Clustering and integration quality control. **a**, UMAPs grouped by sex and treatment group for the acute and repeated datset. **b**, Violin plot showing the number of genes expressed in each cell type. **c**, Violin plot showing the number of unique molecular identifiers (UMIs) within each cell type. **d**, Bargraph showing the percentage of each cell type represented by each of the 8 individual GEM wells used for integration. **e**, Ridge plot for Local Inverse Simpson’s Index (LISI) for each cell type. LISI is a measure of integration sucess with higher values indicating a well-mixed cluster.

**Supplementary Figure 3.**
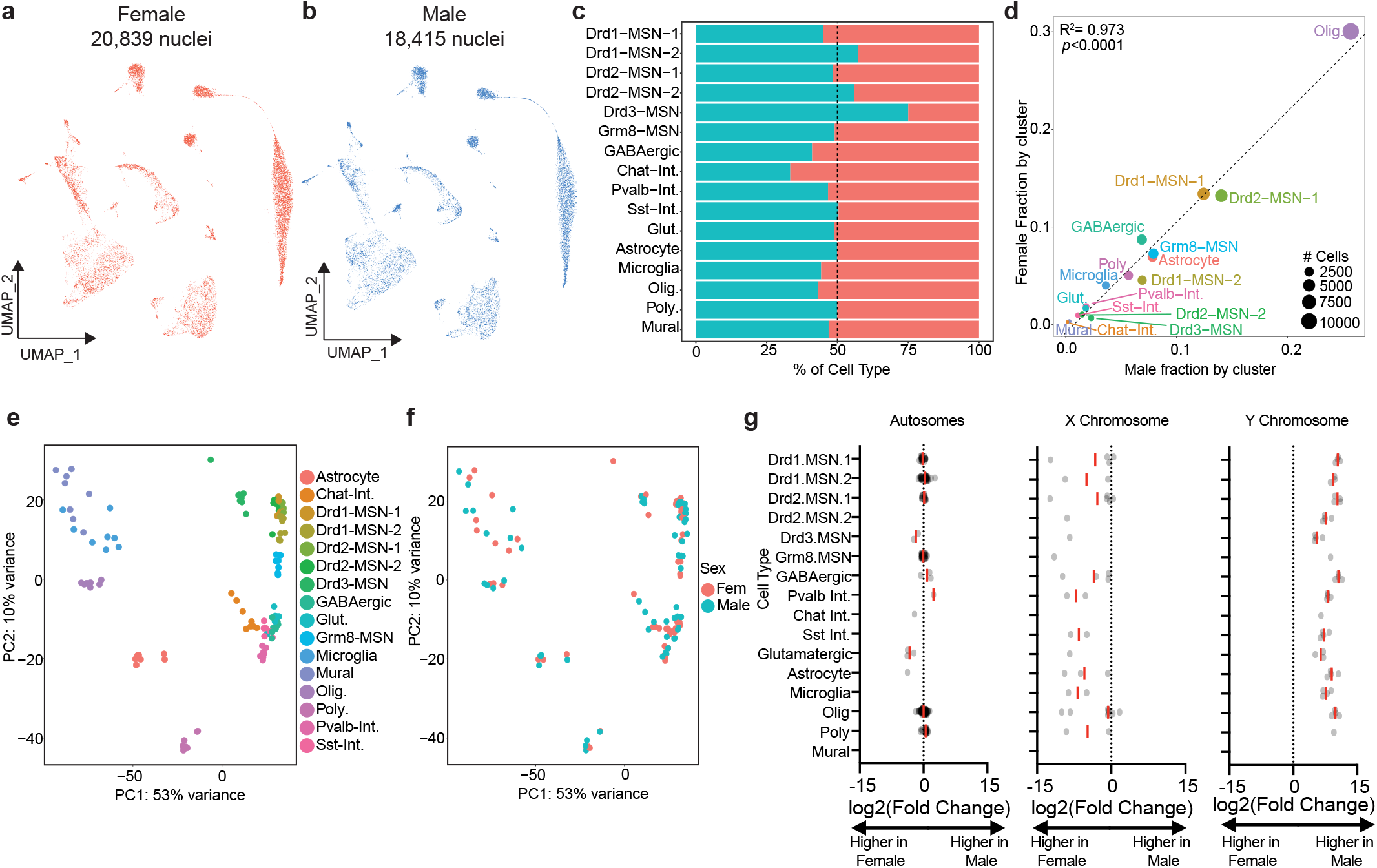
Comparison of baseline transcriptional profiles of male and female cells. **a-b**, UMAPs split by sex. **c**, Percentage of each cell type represented by each sex. **d**, Correlation of the fraction of male and female cells represented by each cluster. Size of the point is representative of the number of cells within the cluster. **e-f**, PCA plots demonstrating that variance within the dataset is largely driven by transcriptional profiles of the cell types and not sex. Each cell type will have 8 individual points that represent 8 GEM wells. **g**, DEGs calculated using a DESeq2-based pseudobulk analysis with dataset and stim used as covariates. Each point on the graph represents a single DEG. The red line represents the average log2(Fold Change) of all DEGs for that cluster.

**Supplementary Figure 4.**
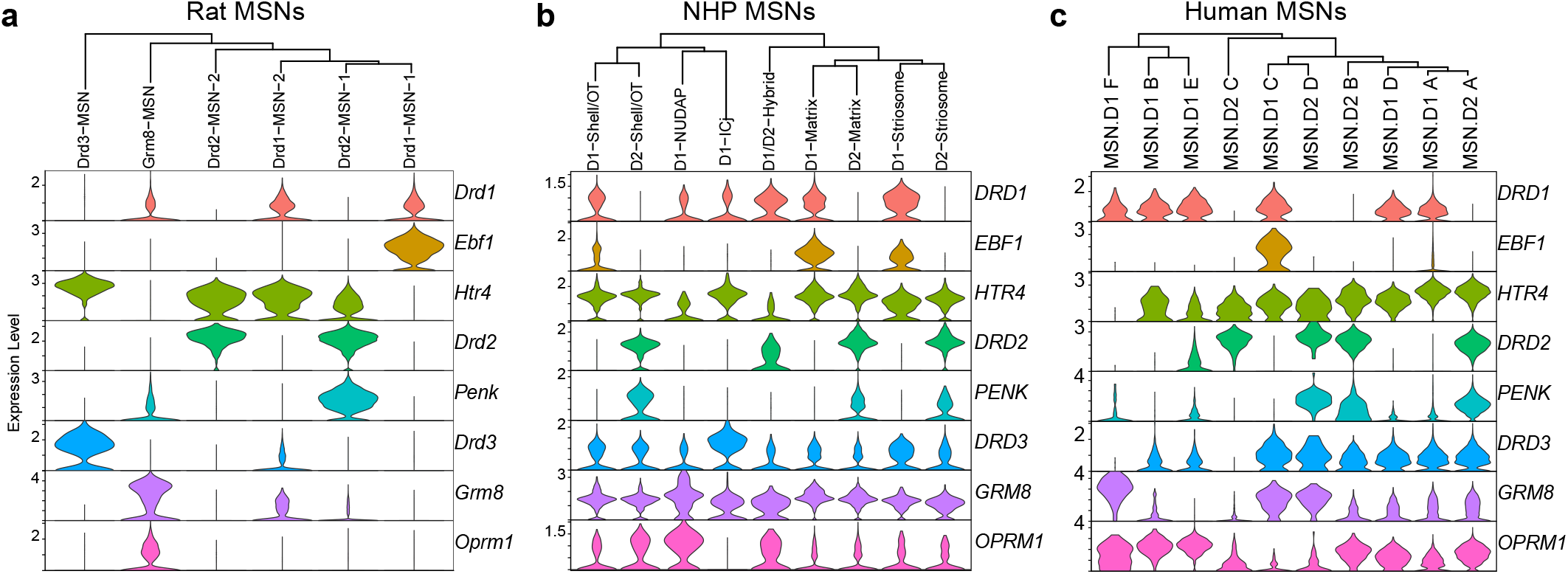
Expression of MSN marker genes across species. **a-c,** Violin plots demonstrating expression of MSN specific markers in rat, NHP, and human MSN clusters.

**Supplementary Figure 5.**
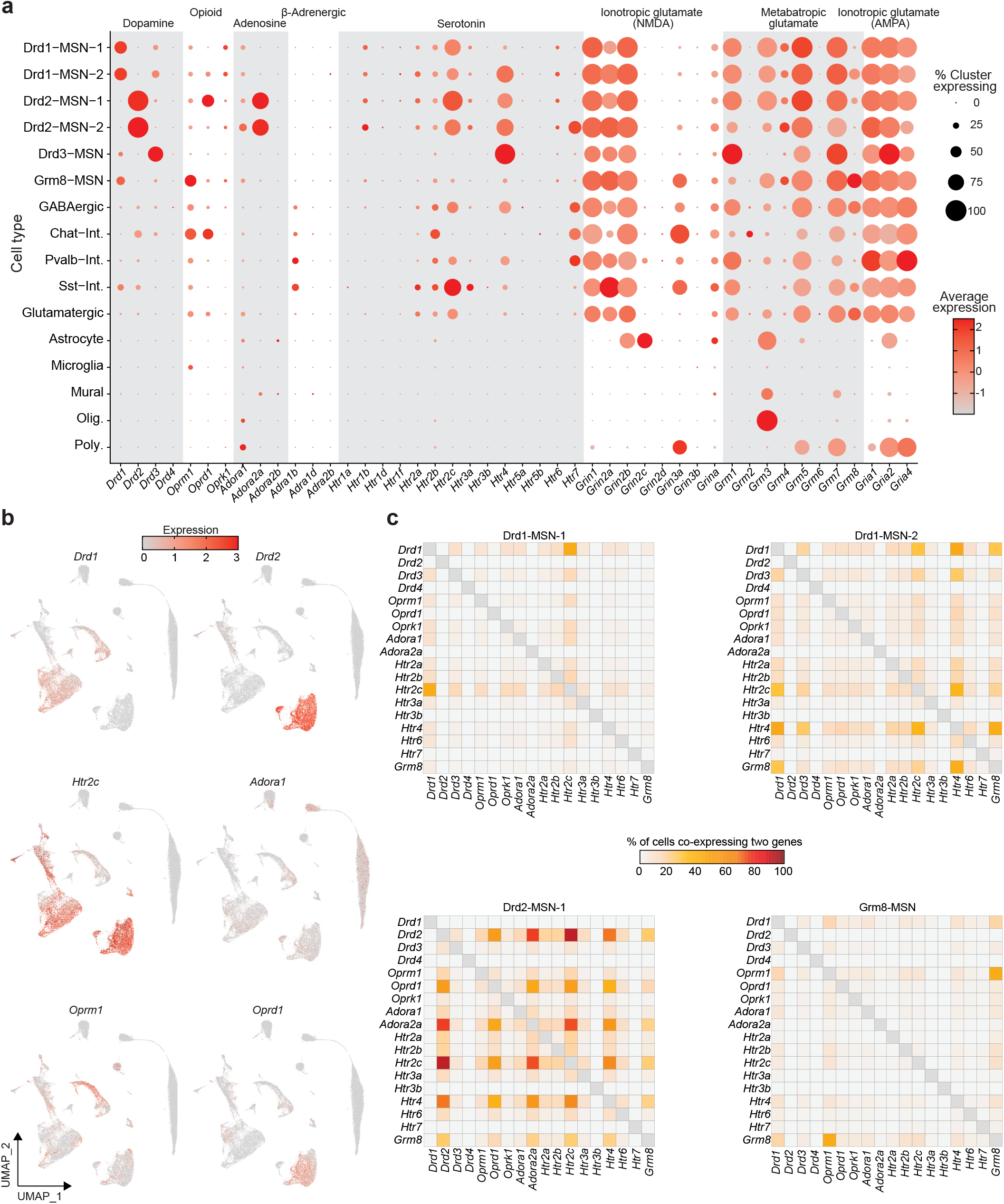
Distribution of expression of genes encoding several receptor systems and subtypes. **a**, Dotplot demonstrating level of expression, and percentage of cells, expressing 46 genes invovled in 8 receptor systems. **b**, UMAPs colored by level of expression of *Drd1*, *Drd2*, *Htr2c*, *Adora1*, *Oprm1*, and *Oprd1*. **c**, Co-expression heatmaps for several receptor systems demonstrating differential expression of serotonic receptors in *Drd1*-expressing MSNs, co-expression of *Drd2* and *Adora2a* in the Drd2-MSN population, and *Grm8* and *Oprm1* in the Grm8-MSN population.

**Supplementary Figure 6.**
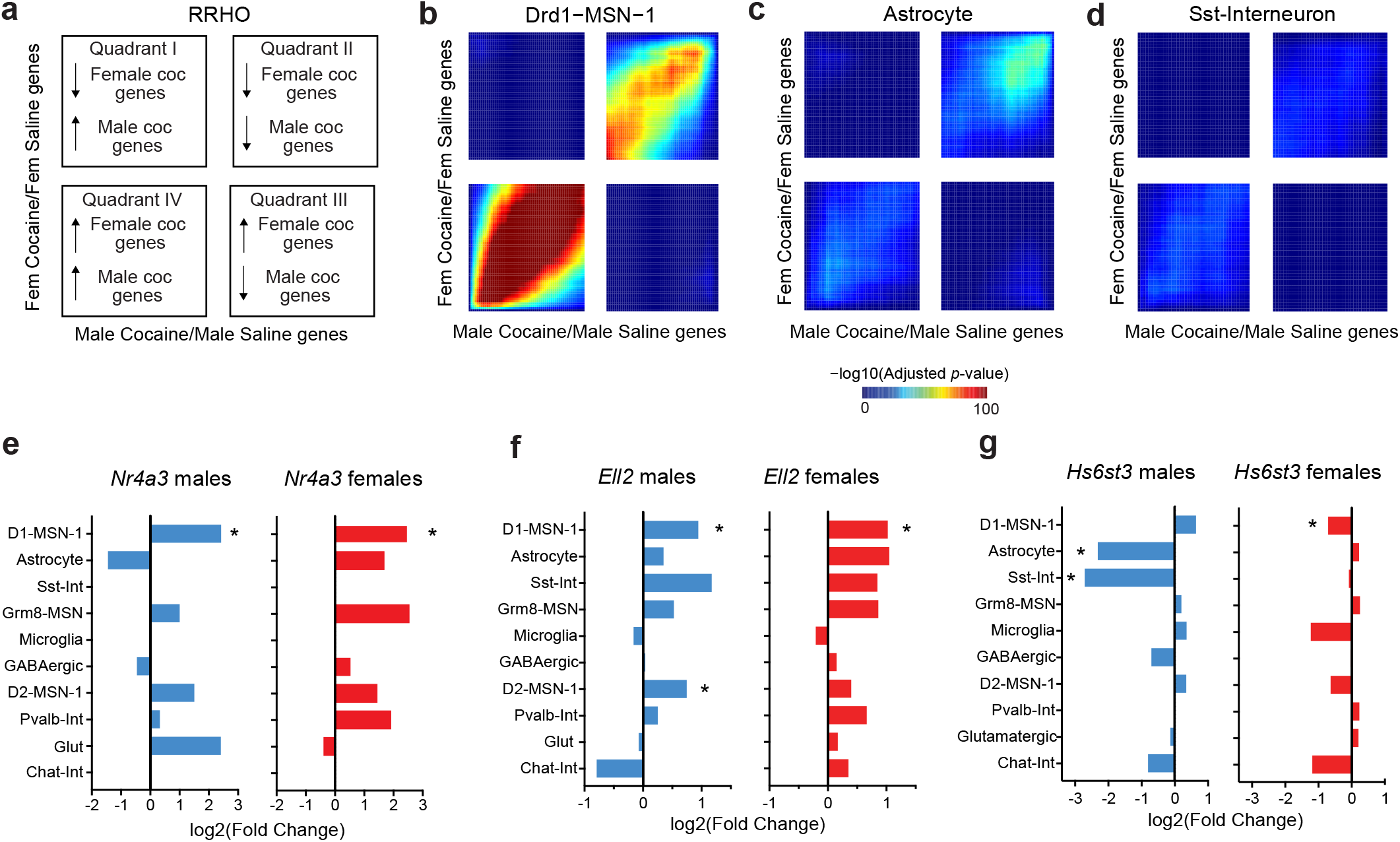
Sex- and cell type-specific comparison of cocaine-induced transcriptional changes identified using pseudobulk differential expression analysis. **a**, Interpretation key for rank-rank hypergeometric overlap (RRHO) heatmaps for sex- and population-specific cocaine genes. Quadrants I and III contain discordant gene changes; quadrants II and IV highlight concordant gene changes. **b-d,** RRHO plots for D1-MSN-1, Astrocyte and Sst-Interneuron populations. Bar graphs illustrating respresentative example genes with similar transcriptional responses to cocaine across populations (**e-f**) and dynamic regulation across populations (**g**).

